# Targeted genome editing in grape using multiple CRISPR-guided editing systems

**DOI:** 10.1101/2022.08.22.504768

**Authors:** Chong Ren, Yanping Lin, Huayang Li, Shaohua Li, Zhenchang Liang

**Affiliations:** Beijing Key Laboratory of Grape Sciences and Enology, Key Laboratory of Plant Resource, Institute of Botany, Chinese Academy of Sciences, Beijing 100093, PR China; University of Chinese Academy of Sciences, Beijing 100049, PR China

**Keywords:** Grape, LbCpf1, xCas9, cytidine base editor, reporter gene

## Abstract

The CRISPR/Cas9 system, together with newly developed CRISPR technologies such as CRISPR/LbCpf1 and base editors, have expanded the scope of targeted genome editing in plants. However, in grape, applications of these novel CRISPR-guided tools have not been reported. Here, we employed EGFP (enhanced green fluorescent protein) and RUBY to help to screen transformed grape cells based on fluorescence and red betalain and tested the activities of CRISPR/LbCpf1, CRISPR/xCas9 and cytidine base editor (CBE) in grape, respectively. The grape *TMT1* (*tonoplastic monosaccharide transporter1*) and *TMT2* genes were simultaneously edited by using LbCpf1, resulting in an efficiency of 16-48%. Furthermore, high temperature (34°C) could enhance editing efficiencies at most of the designed targets. The CRISPR/xCas9 could induce targeted mutagenesis at the target with NGG PAM, but the efficiencies were very low (< 1.9%). The targets with GAA and GAT PAMs that are reported in mammalian cells and rice were not recognized by xCas9 in our study. Moreover, successful C-to-T substitutions were achieved in *GAI1* (*gibberellin insensitive1*) gene by using CBE. The editing efficiencies ranged from 2.4 to 15% at the two targets in *GAI1* in grape cells. Analysis of independent embryos revealed a C-to-T efficiency of 12.5% at the first target of *GAI1*. Taken together, our results demonstrate the efficacy of these CRISPR-guided tools in grape and provide evidence for further application of these editing tools in this economically important species.

## Introduction

The advent of CRISPR (clustered regularly interspaced short palindromic repeats)/Cas9 (CRISPR-associated protein 9) system has revolutionized precise genome editing in mammalian cells and plants. Unlike the other sequence-specific endonucleases ZFNs (zinc-finger nucleases) and TALENs (transcription activator-like effector nucleases), both of which rely on arrays of amino acid modules to bind to specific DNA sequences (Voytas, 2013), Cas9 is an RNA-guided endonuclease that recognizes the target DNA sequence by Watson-Crick base pairing with its single-guide RNA (sgRNA). The protospacer adjacent motifs (PAMs) adjacent to the targets are necessary for target recognition by sgRNA/Cas9 complex (Cong et al., 2013). The commonly used *Streptococcus pyogenes* Cas9 (SpCas9) generally recognizes NGG PAMs, and targets with NAG PAMs were also reported to be cleaved by SpCas9 in rice (Meng et al., 2018). However, the strict PAM requirement when using SpCas9 restricts the range of editable targets. To overcome this limitation, a series of SpCas9 variants have been developed to recognize expanded PAMs. VQR and VRER Cas9 variants could recognize NGA and NGCG PAMs in rice, respectively (Hu et al., 2016). Later, a Cas9 variant named xCas9 was evolved with broad PAM compatibility and could recognize NG, GAA and GAT PAMs in mammalian cells (Hu et al., 2018). In addition, an engineered Cas9 variant, SpCas9-NG, was also reported to recognize relaxed NG PAMs (Nishimasu et al., 2018). The xCas9 and SpCas9-NG variants have been adopted for genome editing in rice to broaden the targeting range (Zhong et al., 2019; Hua et al., 2019; Wang et al., 2019). Recently, SpRY, a near-PAM-less Cas9 variant, was developed to target almost all PAMs in human cells (Walton et al., 2020). CRISPR/SpRY-based tools were accordingly developed for PAM-less genome editing in plants (Xu et al., 2021; Li et al., 2021; Ren et al., 2021a).

In contrast to SpCas9, Cpf1 (also known as Cas12a) recognizes T-rich PAMs, which are located at 5’ of the target sites (Zetsche et al., 2015). Moreover, the single CRISPR RNA (crRNA) for Cpf1 is much shorter (~43 nt) than sgRNA for SpCas9 (Zetsche et al., 2015; Zaidi et al., 2017). More importantly, Cpf1 has RNase III activity and can process premature crRNAs (pre-crRNAs) to produce mature crRNAs (Zaidi et al., 2017; Zetsche et al., 2017), making it an attractive tool for multiplex genome editing with a single crRNA array. Due to high efficiency and specificity, Cpf1 proteins from *Acidaminococcus sp. BV3L6* (AsCpf1), *Francisella novicida* (FnCpf1) and *Lachnospiraceae bacterium ND2006* (LbCpf1) have been used for precise genome editing in plants, including Arabidopsis, rice, soybean, tobacco, citrus and cotton (Endo et al., 2016; Xu et al., 2017; Wang et al., 2017; Kim et al., 2017; Jia et al., 2019; Li et al., 2019).

Mutation in either of the two nuclease domains (D10A mutation in RuvC or H847A mutation in HNH) of Cas9 creates a Cas9 nickase (Cas9n) (Cong et al., 2013; Mali et al., 2013). Cas9n-dependent base editors are recently developed for precise nucleotide substitutions. Cytidine base editor (CBE) was the first developed base editing tool in human cells (Komor et al., 2016), and it consists of a Cas9n (D10A), a cytidine deaminase and an uracil DNA glycosylase (UDG) inhibitor (UGI). The cytidine deaminase could catalyze the conversion of cytosine (C) to uracil (U) in DNA, while the UGI prevents UDG from removing U from DNA in cells (Komor et al., 2016). The reserved U can be further converted into thymine (T) through DNA mismatch repair pathway that is induced by the nick on the target strand generated by Cas9n. Similarly, adenine base editor (ABE) is a fusion of adenosine deaminase and Cas9n, and it enables A-to-G substitutions in cells (Gaudelli et al., 2017). Following the application of CBE and ABE in human cells, base editors were swiftly optimized and applied in plants (Monsur et al., 2020; Molla et al., 2021; Azameti and Dauda, 2021).

The CRISPR/Cas9-mediated genome editing in grape (*Vitis vinifera* L.) was first reported in 2016 (Ren et al., 2016; Malnoy et al., 2016), and an increasing number of reports on successful genome editing using CRISPR/Cas9 have been documented in this species these years (Nakajima et al., 2017; Wang et al., 2018; Ren et al., 2020; Li et al., 2020; Wan et al., 2020; Sunitha and Rock, 2020; Ren et al., 2021b). However, targeted genome editing with newly developed CRISPR-based tools has not been reported in grape to date. In this study, we performed genome editing using the CRISPR/LbCpf1 system in grape by targeting the *TMT* (*tonoplastic monosaccharide transporter*) genes. Furthermore, we tested the activity of xCas9 in recognizing different PAMs in grape. Moreover, base editing using CBE was also conducted. Our results demonstrate the efficacy of these CRISPR-guided tools in grape and provide evidence for further use of these editing tools in this important fruit crop.

## Results

### Evaluation of different selectable markers for selection of transformed cells

To better select transformed cells after transformation, we evaluated four visible markers, namely β-glucuronidase (GUS), EGFP (green fluorescent protein), VvMYBA1 (Ren et al., 2022), and RUBY (He et al., 2020) (Table 1). All the marker genes driven by CaMV35S promoter were introduced into 41B embryogenic cells through *Agrobacterium*-mediated transformation. After transformation, transformed cells were detected at different time points during the screening procedure. The results showed that among the four tested markers, GUS staining was the most sensitive approach to detect transformed cells, given that obvious stained cells were observed at 2 days after transformation (Figure 1). Contrary to GUS, EGFP fluorescence was detected to be very weak at 7 days after selection, and obvious fluorescence was observed 3 months later after selection (Figure 1). Though overexpression of *VvMYBA1* resulted in biosynthesis of red anthocyanin in transformed cells in the presence of light, the phenotype of anthocyanin accumulation, however, was not stable during subcultures.

**Table 1.**
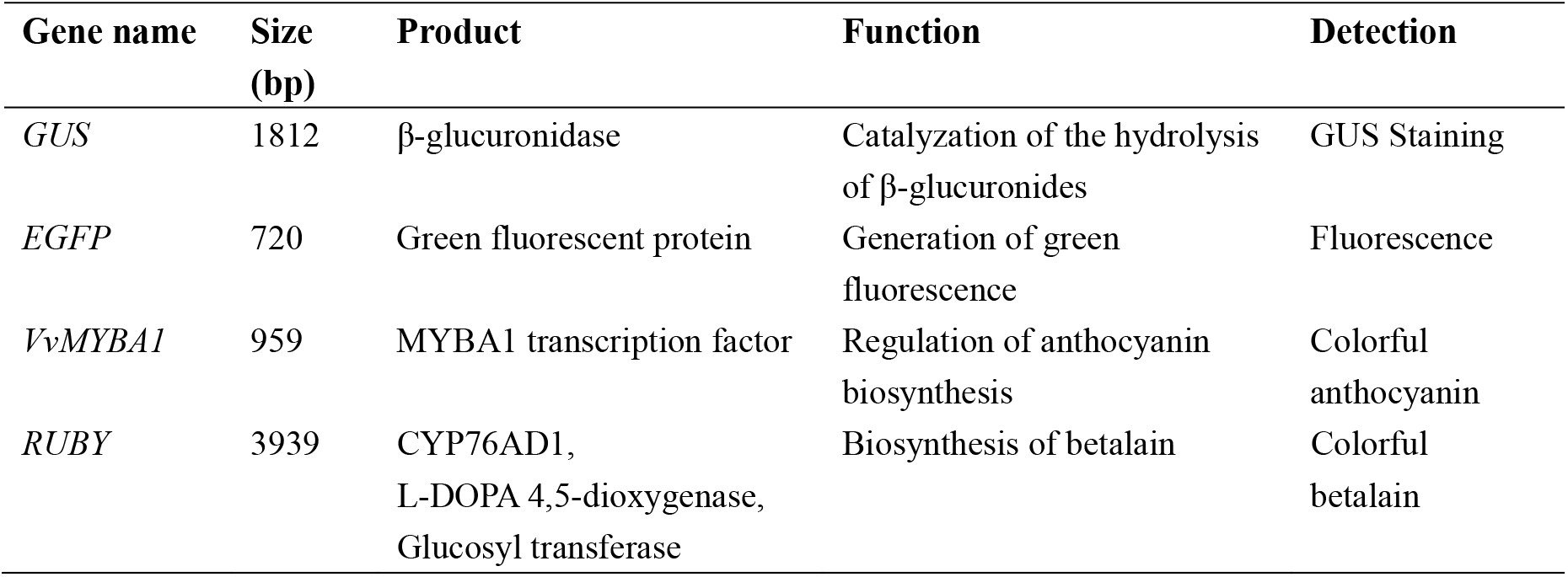
Marker genes tested in this study.

**Figure 1.**
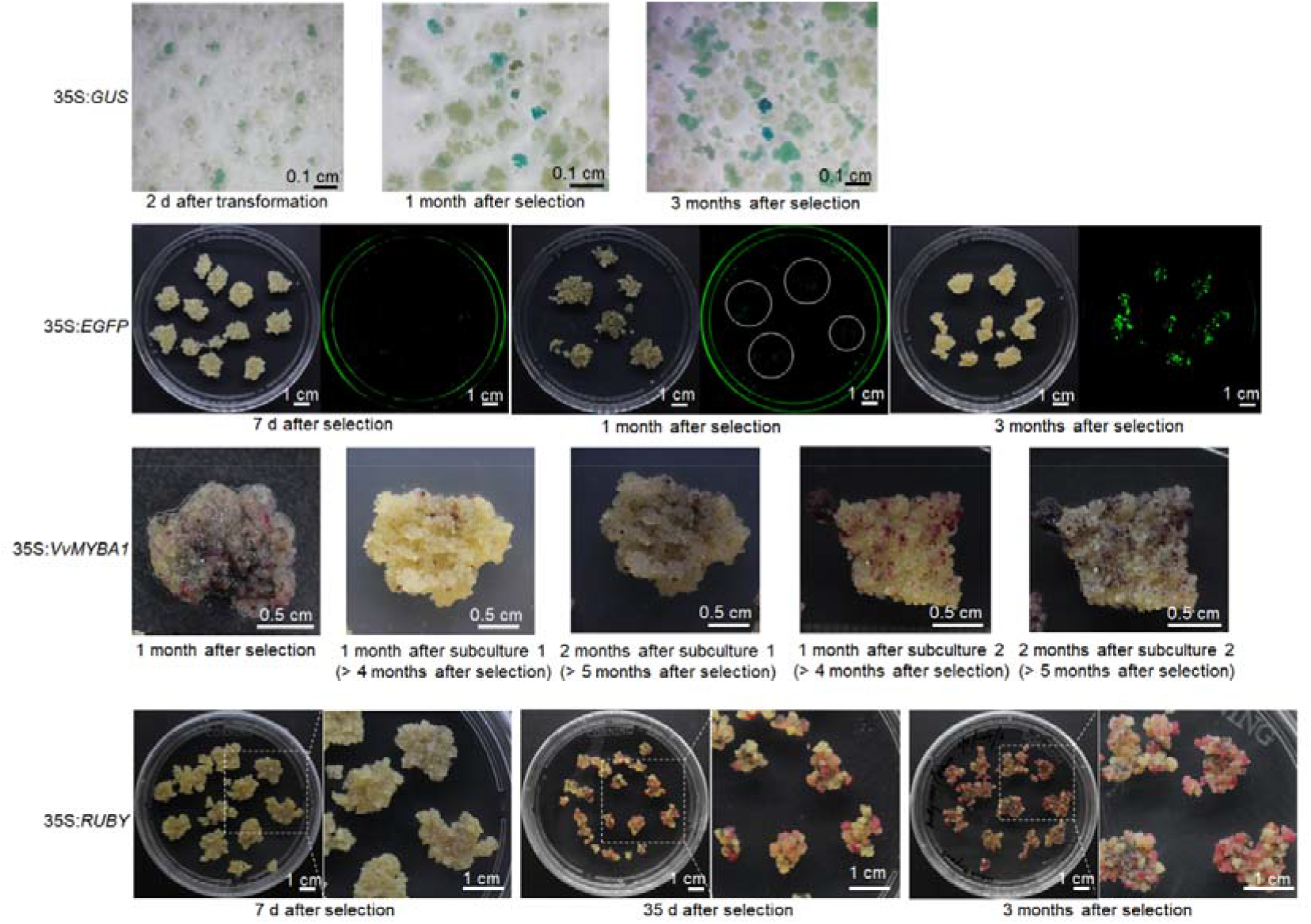
Evaluation of different markers for selection of transformed grape cells. The *GUS* (β-glucuronidase), *EGFP* (enhanced green fluorescent protein), *VvMYBA1*, and *RUBY* driven by CaMV35S (35S) promoter were introduced into 41B embryogenic cells through *Agrobacterium*-mediated transformation, respectively. Grape cells were photographed at different time points as indicated in the figure during the selection.

Grape cells cultured in different flasks (subculture 1 and 2) exhibited different phenotypes in anthocyanin accumulation, and a long-time culture also attenuated the accumulation of anthocyanin in grape cells (Figure 1). In contrast, the use of *RUBY* reporter gene in grape cells resulted in the biosynthesis of vividly red betalain, which is quite stable during subcultures. However, the accumulation of betalain was observed at 35 days in transformed cells after selection (Figure 1). Based on these results, the three visible markers, GUS, EGFP and RUBY, were adopted for visualization of transformed cells during subsequent experiments.

### CRISPR/LbCpf1-mediated targeted mutagenesis of *TMTs*

To test the efficacy of CRISPR/LbCpf1 system in grape, we selected two targets within the exons of *TMT1* and *TMT2* genes (Ren et al., 2021b), respectively (Figure 2A). The corresponding crRNA sequences interspaced by direct repeat (DR) were commercially synthesized and driven by the VvU3.1 promoter (Ren et al., 2021b). The LbCpf1 encoding gene driven by a 35S promoter was ligated into pCAMBIA2300 vector, and the *EGFP* expression cassette was also included to facilitate the selection of transformed cells (Figure 2B). With the aid of EGFP fluorescence, transgenic cells were obtained rapidly after transformation (Figure 2C), which were further identified with exogenous T-DNA insertions by PCR using *Cpf1*-specific primers (Figure 2D). To detect mutations in *TMT1* and *TMT2*, the fragments containing the designed targets were amplified and analyzed by Sanger sequencing. According to the results, nucleotides deletions were detected at the two targets (*TMT1*-crRNA1 and *TMT1*-crRNA2) in *TMT1*, with an editing efficiency of 40%. Of the two targets in *TMT2*, *TMT2*-crRNA1 was detected with nucleotides deletions, with an efficiency of 20% (Figure 2E). These results showed that LbCpf1 could generate targeted mutagenesis in grape, and multiplex genome editing could be achieved using single crRNA array.

**Figure 2.**
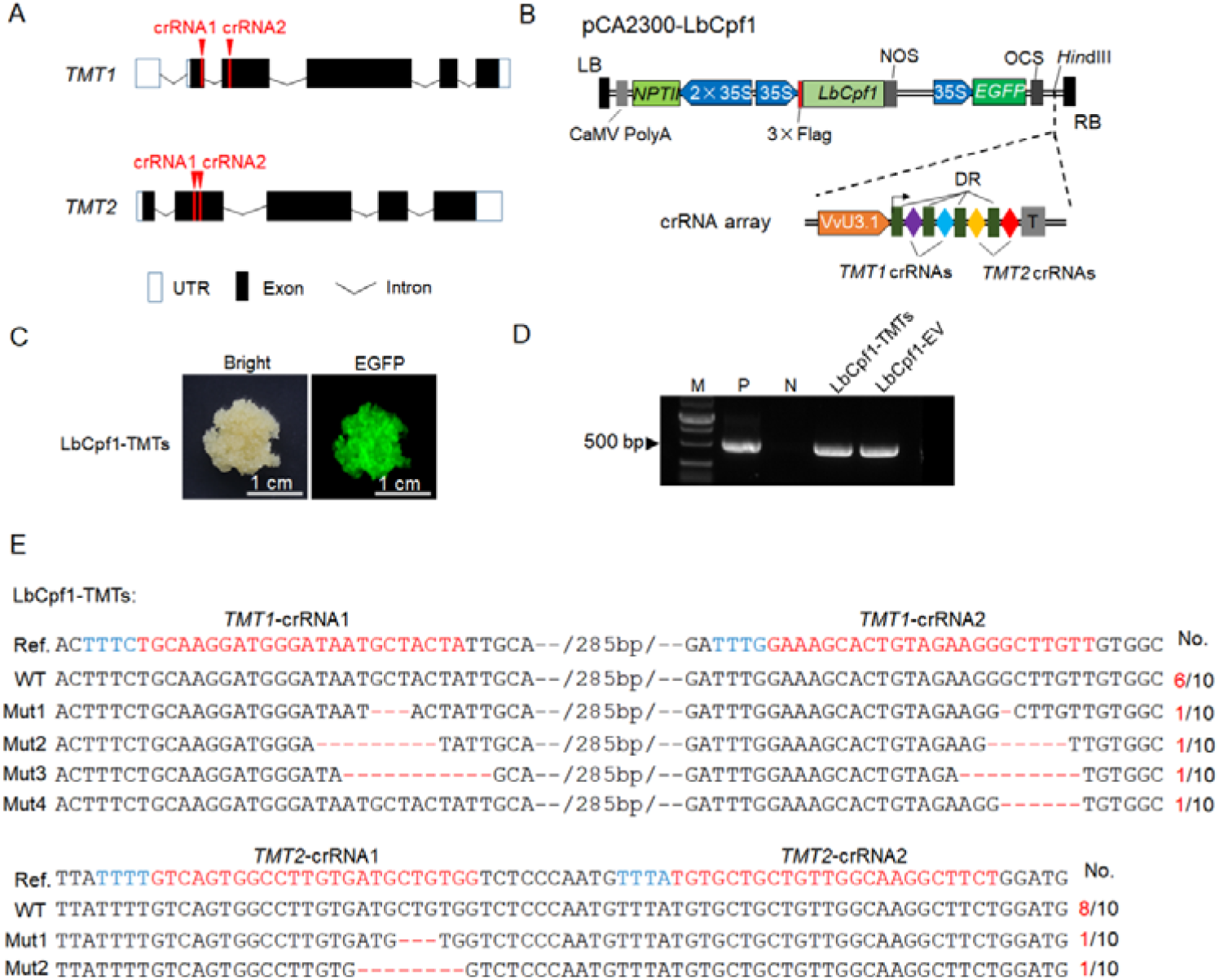
CRISPR/LbCpf1-mediated targeted mutagenesis of the *TMTs*. A, Schematic diagrams of the *TMTs* structure and targets design. The targets within *TMT1* and *TMT2* are indicated by red arrows. B, Schematic illustration of pCA2300-LbCpf1 vector. The *LbCpf1* driven by 35S promoter was cloned into our previously developed pCAMBIA2300-EGFP vector. The designed crRNAs interspaced by mature DRs (direct repeats) were driven by VvU3.1 promoter. The crRNA array was ligated into pCA2300-LbCpf1 via the *Hin*dIII site. *NPTII*, neomycin phosphotransferase gene; NOS, terminator of nopaline synthase gene; OCS, octopine synthase terminator; T, polyT sequence; LB, left border; RB, right border. C, Transgenic cells with EGFP fluorescence. D, Identification of exogenous T-DNA insertion using *Cpf1*-specific primers. The plasmid and wild-type cells were used as positive (P) and negative (N) control, respectively. EV, empty control; M, DNA marker. E, Editing results of the *TMTs*. The target sequences were amplified from transgenic cells shown in C and cloned into pLB vector. A number of 10 amplicons were analyzed by Sanger sequencing for each gene. The target sequences are indicated in red and the PAMs are indicated in blue. Deletions of nucleotides are denoted by red dashed lines, and the number (No.) of mutated (Mut) sequences are shown on the right. Ref, reference sequence; WT, wild-type sequence.

Previous studies revealed that the activity of LbCpf1 was affected by temperatures (Kleinstiver et al., 2019; Li et al., 2021). We then treated the grape cells at 34°C for 7 days, while the cells cultured at 26°C were sampled as the control (Figure 3A). To evaluate the effect of high temperature on genome editing, the target sequences amplified from the control and treated cells were analyzed using Hi-TOM tool (Liu et al., 2019). For *TMT1*, the editing efficiency at 26°C was about 47.7%, and the efficiency detected at 34°C was around 46.5%; for *TMT2*, however, the editing efficiency was increased from 16.1% (26°C) to 24.4% (34°C) (Figure 3B; Supplemental Table S1). Further analyses of individual targets uncovered that the editing of all the four targets in *TMT1* and *TMT2* was affected by high temperature. The editing efficiencies of *TMT1*-crRNA2, *TMT2*-crRNA1 and *TMT2*-crRNA2 were significantly increased at 34°C, while the efficiency of *TMT1*-crRNA1 was decreased (Figure 3C). The percentage of sequences containing mutations in both of the two targets in *TMT1* or *TMT2* was not obviously changed after high temperature treatment (Figure 3C). Moreover, the transgenic cells were used for regeneration (Supplemental Figure S1) and no much difference was observed between the control and treated cells during the regeneration (data not shown). Mutation frequencies were calculated in regenerated plants. Under normal conditions (26°C), over 78% of regenerated plants were identified with mutations in at least one target. However, the percentage was increased to 88% at 34°C (Figure 3D). In addition, the number of plants carrying mutations in *TMT1*-crRNA2, *TMT1*-crRNA2 and *TMT2*-crRNA1 was also increased (Figure 3D). All these results suggested that a short period of high temperature treatment (34°C, 7 days) could improve the editing efficiency generated by LbCpf1.

**Figure 3.**
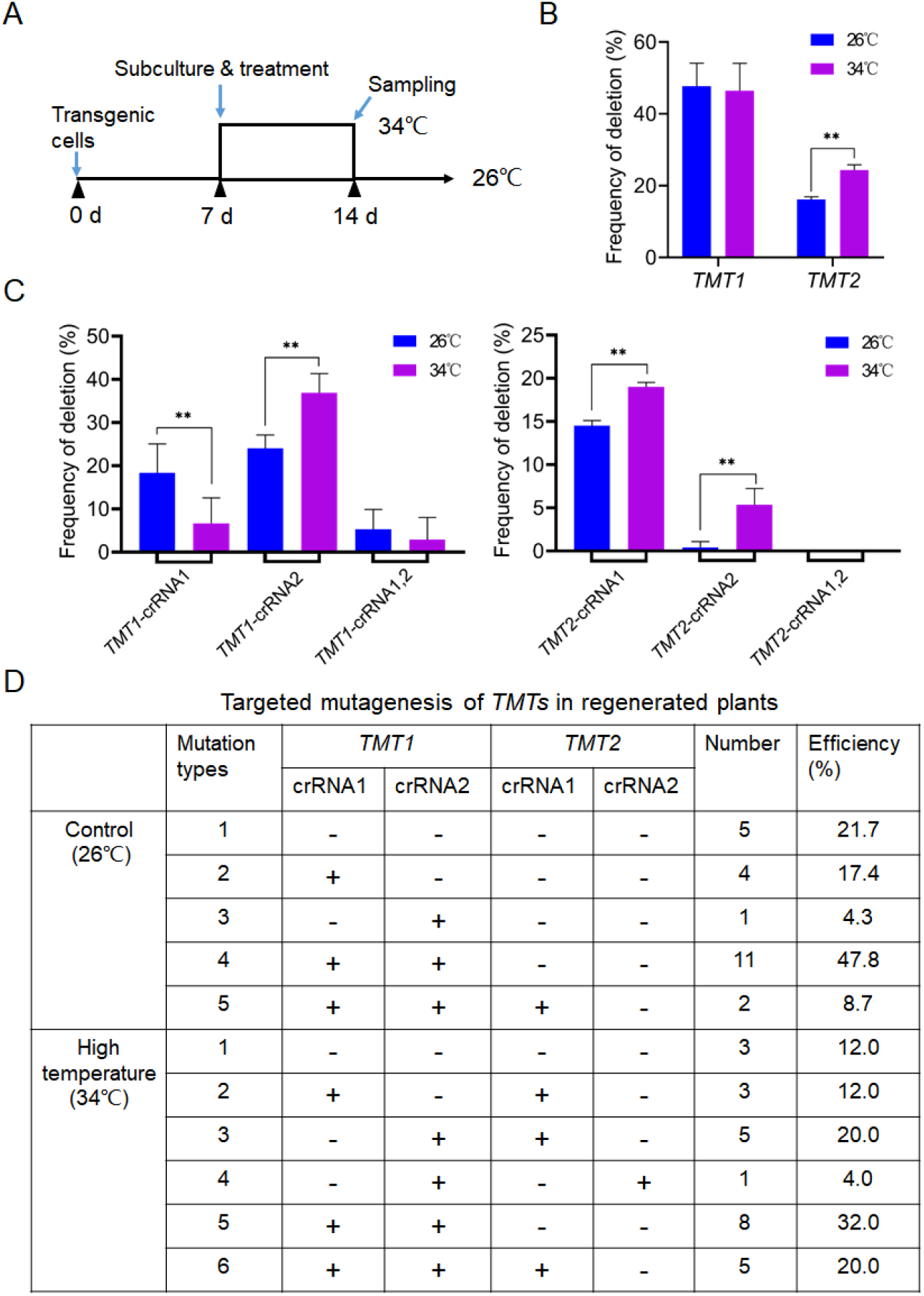
High temperature (34°C) treatment improves activity of LbCpf1 in gene editing. A, Schematic illustration of high temperature treatment. Transgenic cells were subcultured every 7 days at 26°C. To do the treatment, grape cells from the same flask were divided evenly in two, and one flask of cells were shaken at 34°C for high temperature treatment, while the other was incubated at 26°C as the control. Seven days later, both of the cells cultured at 26°C and 34°C were sampled for analysis. B, Editing efficiencies of *TMT1* and *TMT2* under normal and high temperature. C, Effect of high temperature treatment on individual site editing. Data are collected from three replicates and shown as means ± SD. Significant differences were determined using Student’s *t*-test. \*\**P* < 0.01. D, Targeted mutagenesis of the *TMTs* in regenerated plants under normal and high temperature. +, with mutation; −, no mutation.

### CRISPR/xCas9-mediated targeted mutagenesis in grape

To broaden the targeting range in grape genome, we attempted to use xCas9 for genome editing in grape. Given that xCas9 3.7 outperformed xCas9 3.6 in rice (Wang et al., 2019), we therefore employed xCas9 3.7 (hereafter referred to as xCas9) for targeted mutagenesis. To develop the editing construct, the xCas9 sequence was cloned into pCAMBIA1301 vector, just downstream of a 35S promoter. In addition to the intron-containing *GUS* (*iGUS*) gene present in pCAMBIA1301 vector, the *EGFP* expression cassette was also ligated into the vector for selection (Figure 4A). To test the activity of xCas9 in grape genome editing, we designed 12 targets harboring NG, GAT and GAA PAMs (Table 2). Three different genes, *GAI1* (*gibberellin insensitive1*), *TMT1*, and *CCD8* (*carotenoid cleavage dioxygenase8*) (Ren et al., 2020), were involved with each PAM (Table 2). Individual sgRNAs were expressed by VvU6.1 or VvU3.1 promoter (Figure 4A). After *Agrobacterium*-mediated transformation, the grape cells were subject to hygromycin-dependent selection for one month, and the cells survived from the selection (Figure 4B) were checked by GUS staining. The results showed that all the samples except for wild-type (WT) were detected with obvious GUS staining (Figure 4C), suggesting that these cells are successfully transformed. By means of EGFP-assisted selection, the grape cells with strong EGFP fluorescence were enriched (Figure 4D). PCR identification with *xCas9*-specific primers revealed the presence of exogenous T-DNA in these cells (Figure 4E). However, Hi-TOM assay of the target sequences showed that only the first target of *GAI1* (GAI1-g1) with TGG PAM was detected with indel mutations, but the efficiencies observed in two independent experiments were 1.05% and 1.89%, respectively (Figure 4F; Supplemental Table S2). No deletions or insertions were observed in the other targets as expected (Supplemental Table S2). It has been reported that mutation of *GAI1* resulted in gibberellin (GA) insensitive phenotypes (Zhong and Yang, 2011). Considering the low editing efficiencies, we treated the germinated somatic embryos with exogenous GA3, expecting to screen GA3-insensitive plants. Among the 128 analyzed plantlets, two showing relatively short hypocotyls were identified as heterozygous *gai1* mutants (Figure 4G-H). Both of them contained WT and mutated sequences (1-bp insertions) (Figure 4H). All these results showed that xCas9 could work in grape, but the activity was extremely low. Optimization of the CRISPR/xCas9 system should be conducted prior to further application in this species.

**Table 2.**
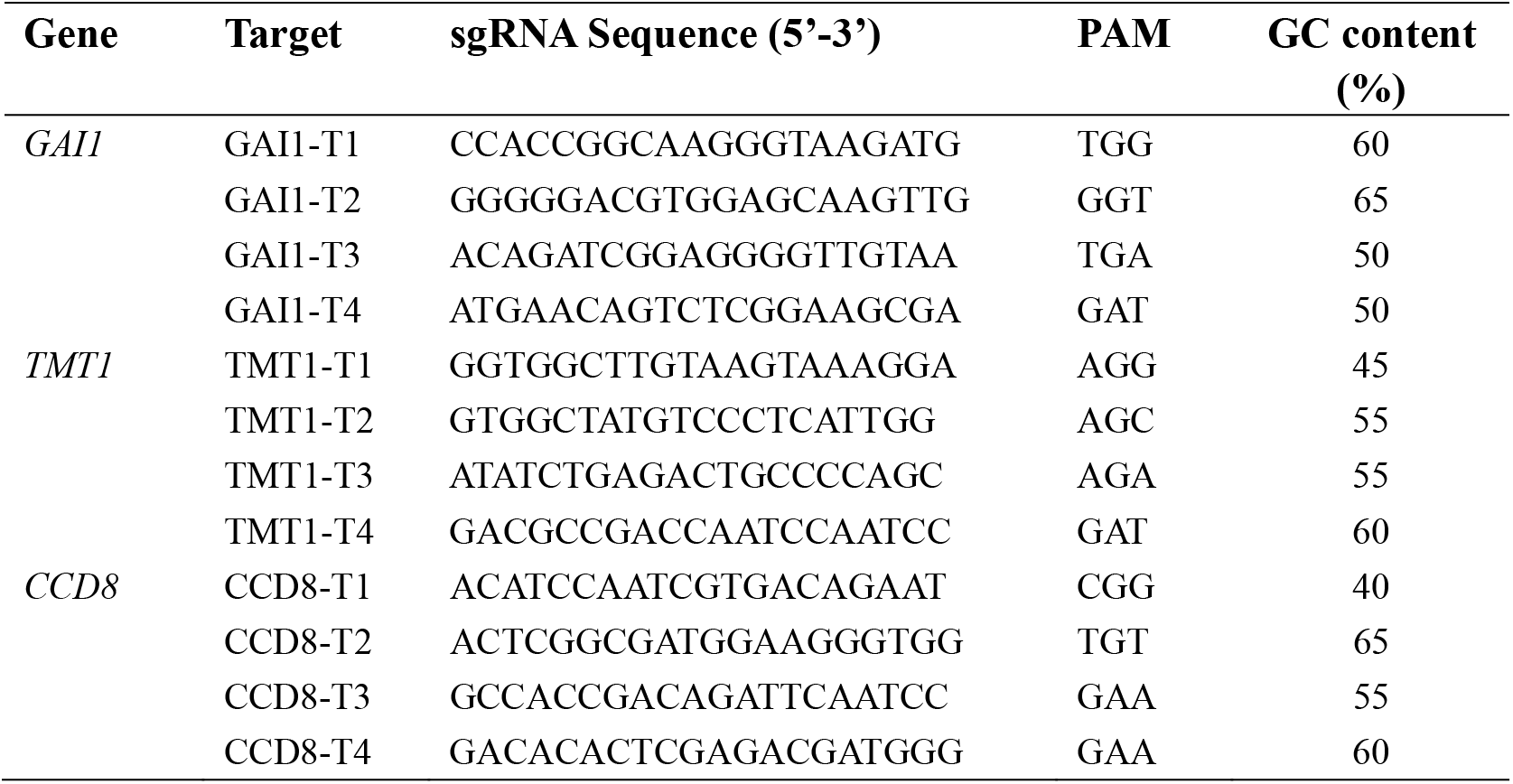
Designed targets for genome editing using xCas9.

**Figure 4.**
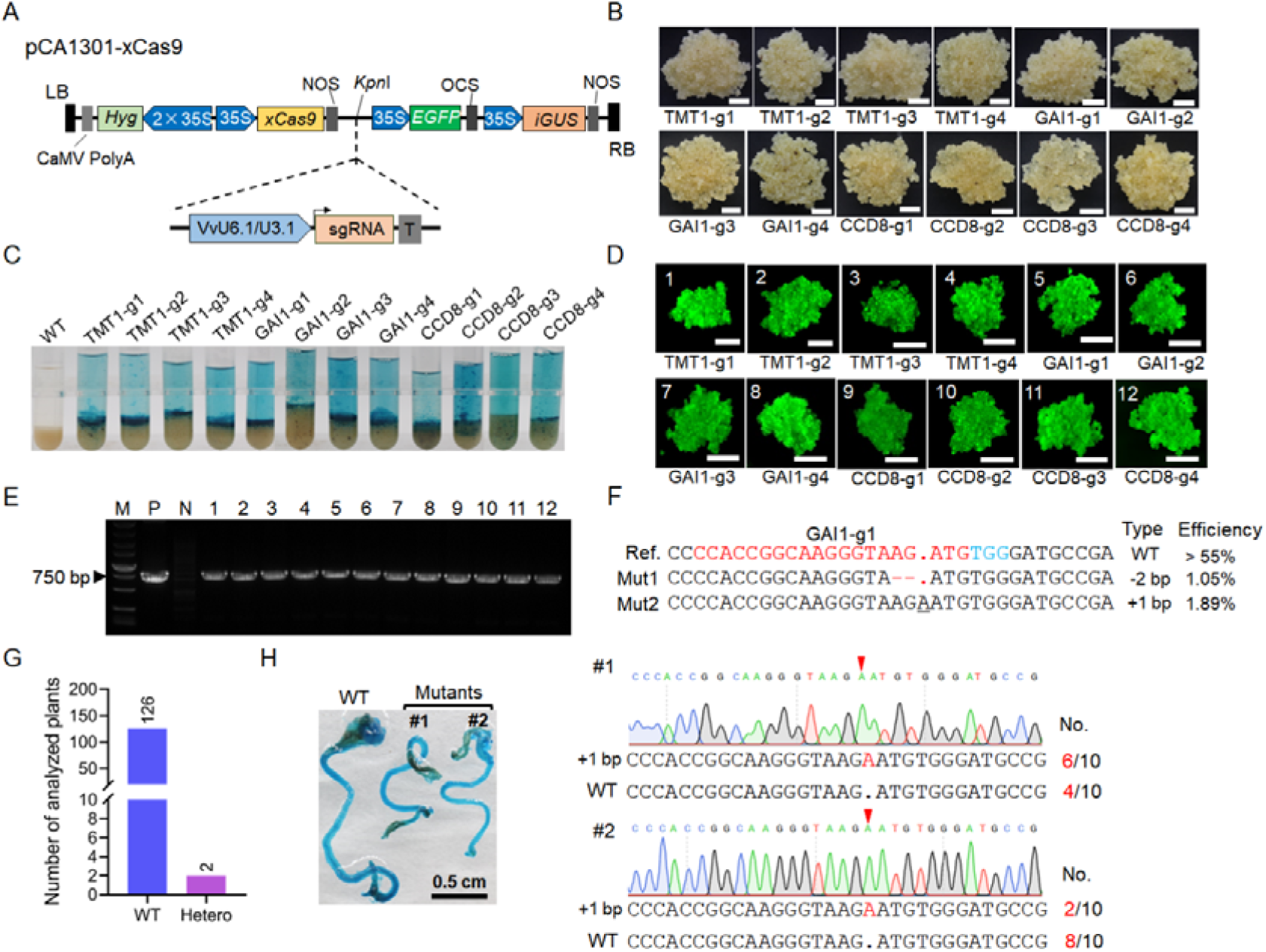
CRISPR/xCas9-mediated targeted mutagenesis of the *GAI1*. A, Schematic illustration of pCA1301-LbCpf1 vector. The *xCas9* driven by 35S promoter was cloned into pCAMBIA1301 vector, which also contains *EGFP* and *GUS* expression cassettes. The sgRNA expression cassette driven by VvU6.1 or VvU3.1 promoter was ligated into pCA1301-LbCpf1 via *Kpn*I site. *Hyg*, hygromycin resistant gene; *iGUS*, intron-containing *GUS* gene. B, Photographs of grape cells after one-month selection. Scale bars: 0.5 cm. C, GUS staining of cells shown in B. The wild-type (WT) cells were shown as negative control. D, EGFP-assisted enrichment of transformed cells. Representative images of grape cells with EGFP signal were shown for each construct. Scale bars: 1 cm. E, PCR identification of exogenous T-DNA insertion using *xCas9*-specific primers. The plasmid and WT cells were used as positive (P) and negative (N) control, respectively. Lines 1-12, the samples shown in D. F, Mutated sequences identified from two replicates. The target sequences were amplified from transgenic cells and subject to Hi-TOM assay. The target sequence and PAM are indicated in red and blue, respectively. The experiment repeated twice. G, Number of analyzed plantlets after regeneration. Hetero, heterozygote. H, Photos of *GAI1* mutants and corresponding mutated sequences. Germinated embryos were cultured on regeneration medium supplemented with 0.01 mg/L GA3 to screen GA3-insensitive mutants. The screened mutants were further confirmed by GUS staining and Sanger sequencing assay. The target sequence was amplified and a number of 10 amplicons were analyzed. The number of analyzed sequences are shown on the right. The insertions are indicated by red arrows.

### Base editing in grape using CBE

To verify the function of CBE in base editing in grape, we first developed the CBE expression construct by modifying pBI121 vector. The BE3 element amplified from the pCMV-BE3 (Komor et al., 2016) was cloned into pBI121 vector, in which the BE3 is driven by a 35S promoter (Figure 5A). Furthermore, the *RUBY* reporter gene driven by a 35S promoter was also ligated into the vector for selection (Figure 5A). Four sgRNAs were designed to test the activities of BE3 in grape. Among the four sgRNAs, two were designed to target the *GAI1* gene, and the other two were used to target *CCD8* and *ALS* (*acetolactate synthase*) gene, respectively (Figure 5B). After RUBY-assisted selection, transformed cells accumulated red betalain were enriched (Figure 5C). PCR identification revealed that these red cells contained exogenous T-DNAs (Figure 5D). Analysis of the target sequences showed that successful C-to-T substitutions were occurred at the two targets (*GAI1*-sgRNA1 and *GAI1*-sgRNA2) in *GAI1* (Figure 5E; Supplemental Table S3), and the editing efficiencies varied from different positions (Figure 5E). The cytidines at position 1-5 (C_1_-C_5_) were primarily edited, with an efficiency being over 5% (Figure 5E). In addition to the C-to-T conversions, other types of substitutions such as G-to-A and A-to-C were also observed within the targets (Supplemental Table S3). By contrast, the target sequences isolated from WT and control cells transformed with *RUBY* reporter gene alone were identified with no C-to-T conversion at the target regions (Supplemental Figure S2, Table S4). However, C-to-T substitutions were detected outside of the targets in both WT and control cells (Supplemental Figure S2, Table S4), indicating the presence of single nucleotide polymorphisms (SNPs). Interestingly, A-to-G and A-to-C substitutions were also detected in control cells (Supplemental Figure S2, Table S4), which suggested that long-time *in vitro* culture or transformation may also lead to nucleotides substitutions in grape cells. Nucleotides deletions were also observed at *GAI1*-sgRNA1 and *GAI1*-sgRNA2 (Figure 5F), but the frequencies were extremely low (< 1.2%) (Figure 5G). Surprisingly, we did not detect desired C-to-T conversions at the target sites in the *ALS* and *CCD8* genes. On the contrary, undesired A-to-C and T-to-C substitutions were detected within the targets in the *ALS* and *CCD8*, respectively, when compared with the wild-type controls (Supplemental Table S5). A low frequency (4.4%) of C-to-T conversions were detected outside of the target in the *ALS* (Supplemental Table S5). Moreover, the other unexpected substitutions might be accumulated during subcultures of grape cells.

**Figure 5.**
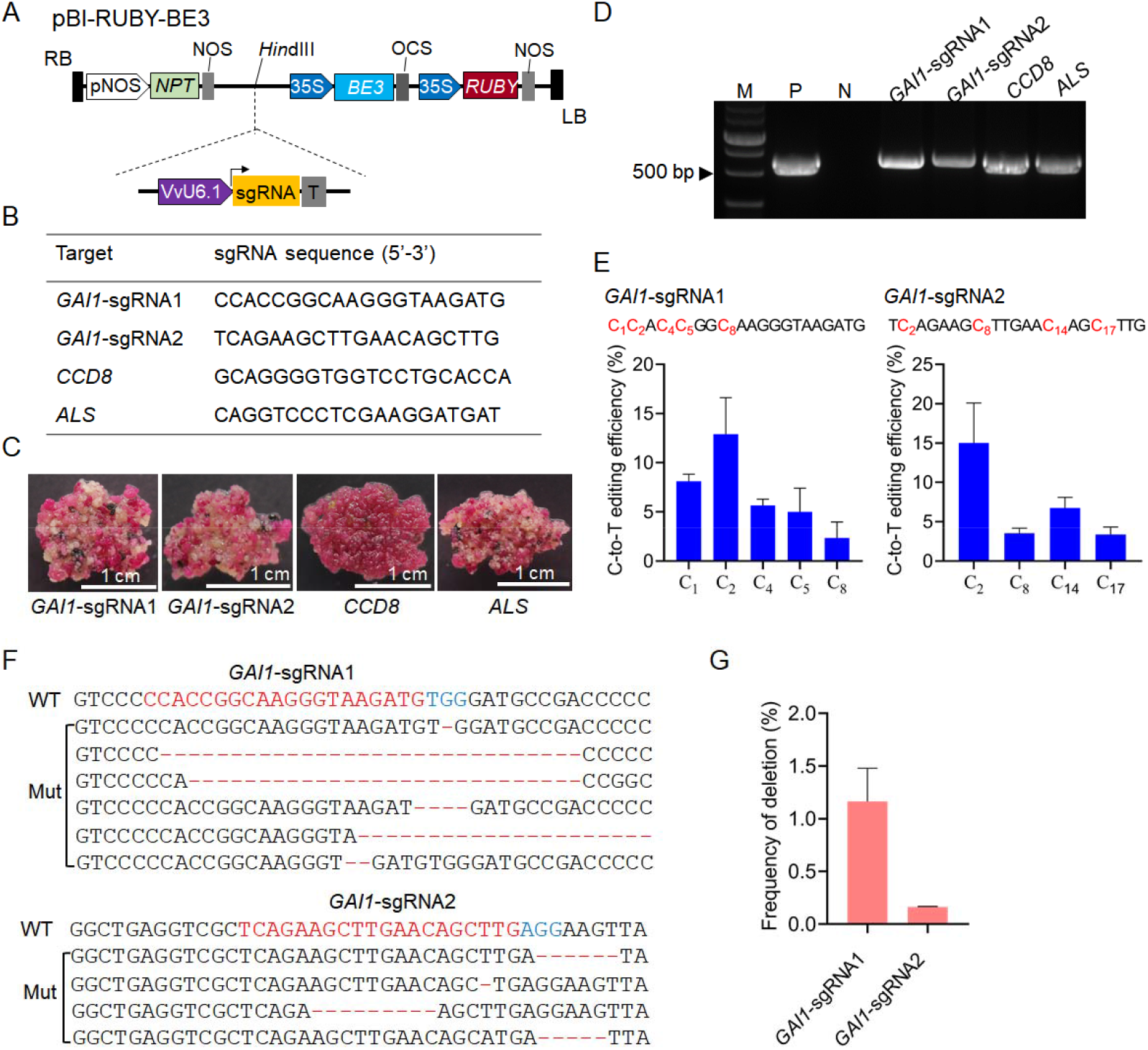
Base editing of the *GAI1* gene using BE3. A, Schematic diagram of pBI-RUBY-BE3 vector. pNOS, promoter of nopaline synthase gene. B, Designed targets for base editing in grape. C, RUBY-assisted enrichment of transformed cells. D, PCR identification of exogenous T-DNA insertion using *BE3*-specific primers. The plasmid and WT cells were used as positive (P) and negative (N) control, respectively. E, C-to-T editing efficiencies within the two targets of the *GAI1*. F, Deletions of nucleotides at the two targets in *GAI1*. G, Frequency of deletions shown in F.

To identify the editing results more accurately, the red cells of *GAI1*-sgRNA1 and *GAI1*-sgRNA2 were used for regeneration. Overexpression of *RUBY* in 41B cells had no much influence on induction of somatic embryos (Figure 6A). Moreover, the induced embryos could further germinate on regeneration medium (Figure 6A).

**Figure 6.**
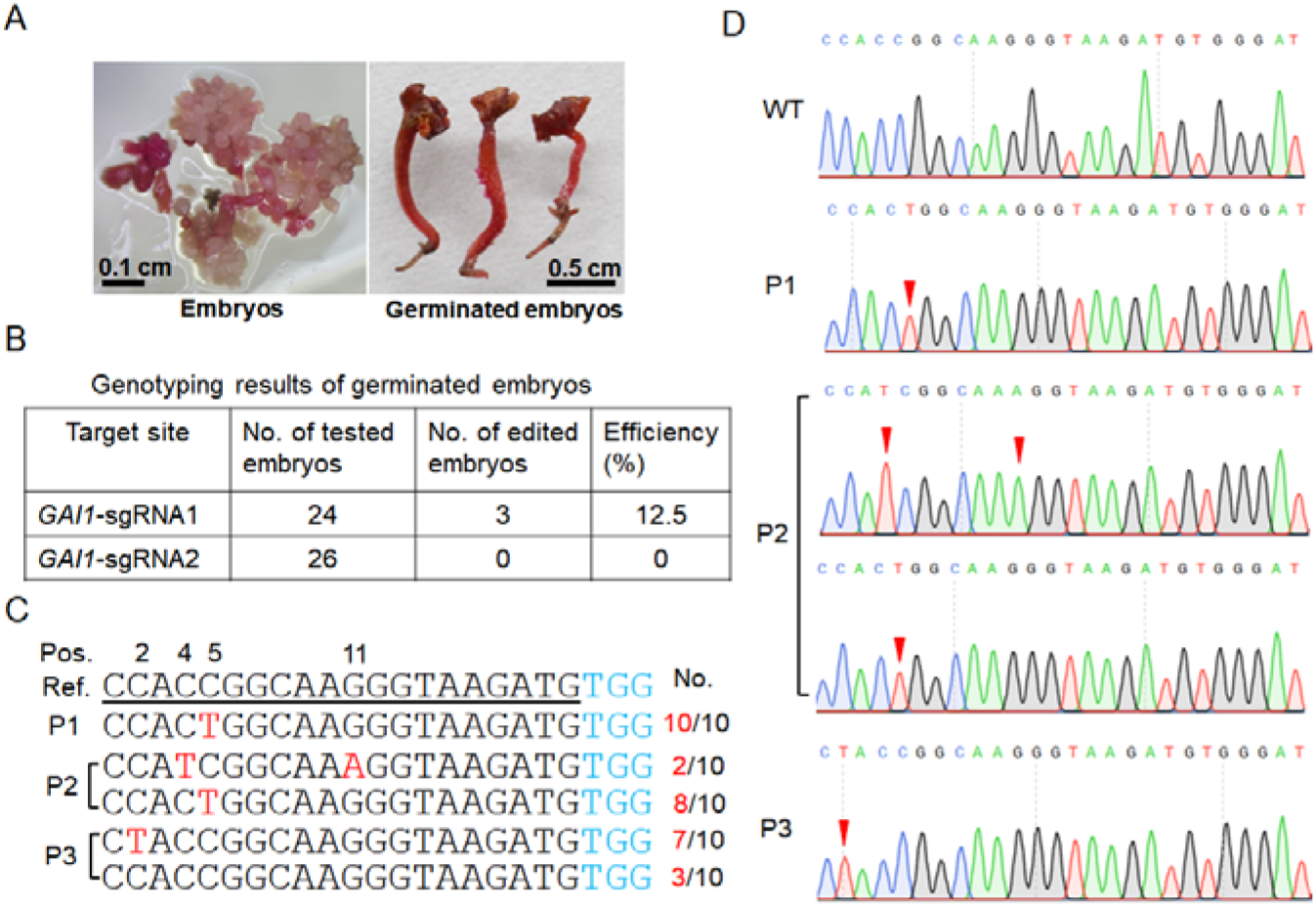
Genotyping results of base editing in germinated embryos. A, Induced and germinated embryos. B, Genotyping results of germinated embryos. C, Mutated sequences identified from the three edited embryos of *GAI1*-sgRNA1. The target sequence is underlined and the PAM is shown in blue. A number of 10 amplicons were analyzed. The mutated nucleotides are indicated in red, and the number of corresponding sequences are shown on the right. Pos, position; Ref, reference sequence; P1-P3: positive embryos. D, Representative chromatograms showing the mutations. The changed nucleotides are denoted by red arrows.

Nevertheless, the germinated embryos could not develop into whole plants (Supplemental Figure S3). Given that RUBY converts tyrosine to betalain in cells (He et al., 2020), we speculated that constitutive consumption of tyrosine may inhibit the process of plant regeneration. Thus, we analyzed the germinated embryos of *GAI1*-sgRNA1 and *GAI1*-sgRNA2, and 3 embryos of *GAI1*-sgRNA1 were found to contain C-to-T conversions (Figure 6B-D). Among the 3 positive embryos (P1-P3), P1 was identified with C-to-T conversion at position 5 (C_5_), while P2 carried C-to-T conversions at position 4 (C_4_) and 5 (C_5_). Additionally, a substitution of G-to-A was also observed in P2 at position 11 (Figure 6C, D). For P3, a C-to-T conversion was detected at position 2 (C_2_) (Figure 6C, D). These results showed that the BE3 could induce desired C-to-T conversions in grape, but undesired mutations may also occur simultaneously.

## Discussion

Grape cells derived from embryogenic callus have been widely used as materials for transformation (Lecourieux et al., 2010; Wang et al., 2021; Ren et al., 2021b). Application of visible markers during selection would be helpful to enrich transformed cells, especially when the transformation rate is low. In this study, we evaluated four different markers, and GUS turned out to be the most sensitive one for the detection of transformed cells (Figure 1). However, GUS staining is destructive to grape cells, so it could be used with the other markers. In contrast, EGFP is not harmful to grape cells, but the signal of EGFP was not easy to be detected at the early stage of selection (Figure 1). Overexpression of *VvMYBA1* and *RUBY* resulted in red colors of grape cells, but the phenotypes generated by RUBY are more stable than that caused by VvMYBA1 (Figure 1). Notably, constitutive expression of *RUBY* in grape cells may hinder plant regeneration, at least in the 41B cells tested in our study (Supplemental Figure S3). To gain more insights into the use of this marker, RUBY could be tested in other cultivars with established transformation systems, such as Chardonnay and Thompson Seedless (Ren et al., 2016; Wang et al., 2018; Li et al., 2020).

With the aid of EGFP marker, we tested the CRISPR/LbCpf1 system in grape genome editing. The editing results showed that LbCpf1 is effective in generating targeted mutagenesis in grape, and is also capable of editing multiple targets simultaneously by using a single crRNA array (Figure 2E). Intriguingly, among the four designed targets, the editing of *TMT2*-crRNA2 was less efficient than the others (Figure 2E, 3C), probably because that the retained polyT sequence attenuated the activity of crRNA2 of *TMT2*. However, high temperature (34°C) treatment could improve the editing efficiency, and the editing efficiency observed for *TMT2*-crRNA2 in grape cells was increased from 0.4 to 5.4% after treatment (Figure 3C). Moreover, the percentage of edited plants was also increased after high temperature treatment (Figure 3D). These results suggest that elevated temperature could improve the activity of CRISPR/LbCpf1 in grape, which is consistent with previous results (Kleinstiver et al., 2019; Li et al., 2021). More recently, temperature-tolerant LbCpf1 has been developed with improved activity for efficient gene editing in plants (Schindele and Puchta, 2020; Huang et al., 2021), which provides an alternative to genome editing using LbCpf1 in grape.

xCas9 is a variant that can recognize expanded PAMs. In mammalian cells, xCas9 was reported to recognize NG, GAA and GAT PAMs (Hu et al., 2018). In our study, we tested cleavage activity of xCas9 on targets with NG, GAA and GAT PAMs (Table 2). However, only the target with NGG PAM (GAI1-g1) was successfully edited, and the efficiencies turned out to be very low (< 1.9%) (Figure 4). Though the CRISPR/xCas9 system could work in grape, the editing efficiency is not acceptable for practical use. In rice, the CRISPR-guided xCas9 is also inefficient for editing targets with GAA and GAT PAMs (Zhong et al., 2019; Hua et al., 2019; Wang et al., 2019). Luckily, the use of tRNA and enhanced sgRNA was reported to be an efficient way to improve the CRISPR/xCas9 system, enabling the system to edit the targets with NG, GAA, GAT, and even GAG PAMs (Zhang et al., 2020). This study provides a promising strategy for optimizing the current CRISPR/xCas9 system used in our study. Moreover, the other Cas9 variants such as Cas9-NG and SpRY could be tested in grape for targeted mutagenesis with relaxed PAMs in the future.

Base editing using BE3 was achieved by targeting the *GAI1* gene in grape (Figure 5). Desired C-to-T conversions were observed in both grape cells and independent embryos (Figure 5 and 6). In grape cells, the main editing window ranged from position 1 to 5 (efficiency ≥ 5%). However, C-to-T conversion was also observed at C_17_ within the *GAI1*-sgRNA2, with a frequency of 6.7% (Figure 5E). Analysis of independent embryos revealed that C_2_, C_4_ and C_5_ could be successfully edited (Figure 6C, D). To define the editing window more precisely, plenty of plants involved with different targets should be analyzed in the future. Notably, SNPs (C/T) were observed in the *GAI* gene in WT 41B cells (Supplemental Figure S2), and similar results were also detected in the *ALS* and *CCD8* genes (Supplemental Table S5). Therefore, it is better to avoid the regions with SNPs in the genes of interest when using BE3. Moreover, to reduce the accumulation of mutations occurred during long-time *in vitro* culture of grape cells, newly induced embryogenic callus could be used for transformation. Recent studies uncovered that CBE induced genome-wide off-target mutations in mouse and rice (Zuo et al., 2019; Jin et al., 2019), whereas ABE exhibited high specificity in base editing (Jin et al., 2019). According to our results, A-to-G conversions happened less frequently in grape cells (Supplemental Table S2, S4-S5). Alternatively, ABE could serve as a potential approach for grape base editing. In addition, the newly developed prime editor that has the ability to accomplish precise base-edits (Anzalone et al., 2019) could also be a promising approach for base editing in grape.

In summary, our results provide evidence for application of CRISPR-guided LbCpf1, xCas9 and CBE in grape. The concerns about the three CRISPR-based systems could be addressed by optimization of these systems in the future. More importantly, the use of these technologies in grape would expand the CRISPR toolbox and facilitate basic research and breeding in this important species.

## Materials and methods

### Plant material, transformation, and regeneration

The 41B (*V. vinifera* × *V. berlandieri*) grape cells were subcultured weekly in glycerol-maltose (GM) liquid medium supplemented with 1 mg/L naphthoxy acetic acid (NOA). The grape cells were transformed via *Agrobacterium* (strain EHA105)-mediated transformation. After transformation, the 41B cells were cultured in liquid GM medium supplemented with 200 mg/L timentin and 5 mg/L hygromycin or paromomycin for selection of transformed cells. To enrich the transformed cells, the grape cells were divided into small cell mass (CM) before subculture, and those CMs with obvious EGFP fluorescence or red cells were manually picked and pooled together in a flask for subculture. For plant regeneration, the screened cells were transferred to regeneration medium (GM without NOA) for induction of embryos. Whole plants were recovered with roots in McCown woody plant medium (Duchefa Biochemie) supplemented with 0.2 mg/L naphthylacetic acid (NAA). Both transformation and regeneration were conducted as previously described (Ren et al., 2020).

### Construction of expression vectors

The well-constructed vectors pBI121-GUS (Ren et al., 2021b), pCAMBIA2300-EGFP (Wang et al., 2021), pCAMBIA2300-VvMYBA1 (Ren et al., 2022) were directly used for transformation of grape cells. To develop the *RUBY* expression vector, the coding sequence of *RUBY* gene was amplified from the pDR5-RUBY vector (He et al., 2020) using the primers RUBY-PCR-F/R (Supplemental Table S6). Then the amplified *RUBY* gene was cloned into pBI121-GUS in place of the *GUS* gene through *Bam*HI and *Sac*I sites via homologous recombination (HR) by using the ClonExpress II One Step Cloning Kit (Vazyme) to develop the pBI-RUBY vector.

To develop the LbCpf1 editing vector, the *LbCpf1* gene was amplified from pCAMBIA-LbCpf1 construct (Wang et al., 2017) by PCR using the primers Cpf1-PCR-F/R. The amplified sequence was then ligated into the modified pCAMBIA2300-EGFP vector, in which the *LbCpf1* gene was driven by the 35S promoter, through *Eco*RI and *Kpn*I sites via HR. The constructed vector was named pCA2300-LbCpf1. The editing targets were designed with no potential off-targets using the CRISPR-GE online tool (http://skl.scau.edu.cn/), and the crRNAs expression array containing the VvU3.1 promoter (Ren et al., 2021b), the designed crRNAs, and mature DRs (Supplemental Figure S4) was commercially synthesized (Tsingke) and ligated into the pCA2300-LbCpf1 vector through *Hin*dIII site using the T4 DNA ligase kit (NEB).

To develop the xCas9 editing vectors, the *xCas9* gene was amplified from the xCas9 3.7 plasmid (Addgene: #108379) by PCR using the primers xCas9-PCR-F/R. The amplified *xCas9* fragment was inserted into the modified pCAMBIA1301 vector through *Eco*RI and *Bam*HI sites via HR. The 35S-EGFP-OCS expression cassette was amplified from pCA2300-LbCpf1 vector using 35EGFP-PCR-F/R primers and ligated into the pCAMBIA1301-xCas9 vector through *Xba*I site via HR. The well-constructed vector was designated pCA1301-xCas9. The sgRNAs with high specificity were designed using the CRISPR-GE tool, and the development of sgRNA expression cassettes were performed as previously described (Ren et al., 2021b). The developed sgRNA cassettes were ligated into pCA1301-xCas9 through *Kpn*I site via HR.

To construct the CBE vector, we amplified the BE3 fragment from the pCMV-BE3 plasmid (Addgene: #73021) using the primers BE3-PCR-F/R. A 35S promoter and OCS terminator were previously cloned into the pBI-RUBY vector, and the amplified BE3 was ligated into the vector just between the 35S and OCS elements through *Hin*dIII site via HR. Similarly, the development of sgRNA expression cassettes were performed as previously described (Ren et al., 2021b). The developed sgRNAs cassettes were ligated into pBI-RUBY-BE3 vector through *Sca*I site via HR. All the primers are available in Supplemental Table S6.

### Identification of exogenous T-DNA insertions

Genomic DNA was extracted using plant genomic DNA extraction kit (Aidlab) following the manufacturer’s instructions. The prepared genomic DNA was used for PCR identification with *Cpf1*-, *xCas9*-, or *BE3*-specific primers (Supplemental Table S6) using Rapid Taq Master Mix (Vazyme) according to the manufacturer’s protocol. The PCR products were separated by 1% agarose gel electrophoresis.

### Mutation detection

Genomic DNA isolated from grape cells, germinated embryos or plants was used as the template for PCR amplification of target fragments. The mutations in target sequences were analyzed by Sanger sequencing or Hi-TOM assay (http://www.hi-tom.net/hi-tom/). For Hi-TOM assay, target-specific primers with bridging sequences at 5’ end (Supplemental Table S6) were used to amplify the target fragments using Rapid Taq Master Mix. The PCR products were used for next-generation sequencing after a second-round PCR called barcoding PCR (Liu et al., 2019). More than 5,000 reads were generated and analyzed for each sample. The assay of mutated sequences was conducted as previously described (Liu et al., 2019). For Sanger sequencing assay, the amplified fragments were ligated into pLB vector (TIANGEN), and a number of 10 amplicons were analyzed for each sample.

### Treatment of grape cells and embryos

For high temperature treatment, the grape cells from one 250 mL flask were subcultured evenly into two 100 mL flasks fitted with 30 mL of liquid GM medium. One flask was shaken at 120 rpm at 26°C in the dark, while the other was shaken at 34°C for high temperature treatment. Seven days later, the cells were collected and used for Hi-TOM assay. Three replicates were conducted for the treatment.

For GA treatment, the germinated embryos were transferred to solid regeneration medium (GM without NOA) supplemented with 0.01 mg/L GA3, and were cultured at 26°C under light condition for 4 weeks.

### GUS staining and microscopy

The x-Gluc was used as the substrate of β-glucuronidase. GUS staining was conducted using the GUS staining kit (Coolaber) according to the manufacturer’s protocol. The stained cells, as well as embryos, were photographed using a stereomicroscope.

## Supplemental data

**Supplemental Figure S1**. Regeneration of plants from grape cells transformed with pCA2300-LbCpf1 construct.

**Supplemental Figure S2**. The target fragment of the *GAI1* gene.

**Supplemental Figure S3**. Regeneration of plants from grape cells transformed with pBI-RUBY-BE3 construct.

**Supplemental Figure S4**. The sequence of crRNA array for LbCpf1-mediated genome editing.

**Supplemental Table S1**. Hi-Tom assay of *TMTs* editing using LbCpf1 under different temperatures.

**Supplemental Table S2**. Editing results of the *TMT1*, *CCD8* and *GAI1* targets using CRISPR-xCas9 system.

**Supplemental Table S3**. Base editing results of the *GAI1* gene.

**Supplemental Table S4**. Hi-Tom assay of the *GAI1* targets in control cells.

**Supplemental Table S5**. Base editing results of the *ALS* and *CCD8* genes.

**Supplemental Table S6**. Primers used in this study.

## Acknowledgments

We thank professor Jian-Kang Zhu (Shanghai Center for Plant Stress Biology and Center for Excellence in Molecular Plant Sciences, Chinese Academy of Sciences) for kindly sharing pCAMBIA-LbCpf1 plasmid, professor Yubing He (Nanjing Agricultural University) for kindly sharing pDR5-RUBY plasmid, and professor Lanqin Xia for kindly sharing pCMV-BE3 plasmid.

